# Cultured deep-sea PVC bacteria shed light on eukaryogenesis

**DOI:** 10.1101/2021.11.19.469327

**Authors:** Rikuan Zheng, Chong Wang, Tianhang Zhang, Yingqi Tan, Chaomin Sun

**Affiliations:** CAS and Shandong Province Key Laboratory of Experimental Marine Biology & Center of Deep Sea Research, Institute of Oceanology, Chinese Academy of Sciences, Qingdao, China; Laboratory for Marine Biology and Biotechnology, Pilot National Laboratory for Marine Science and Technology, Qingdao, China; College of Earth Science, University of Chinese Academy of Sciences, Beijing, China; Center of Ocean Mega-Science, Chinese Academy of Sciences, Qingdao, China; College of Life Sciences, Qingdao University, Qingdao, China

**Author notes:** Corresponding author Chaomin Sun.

**Keywords:** PVC superphylum, eukaryogenesis, nucleus, nucleolus, mitosis

## Abstract

Evolutionary relationship between prokaryotes and eukaryotes continues to fascinate biologists. Accumulated studies suggest a eukaryogenesis model based on the PVC (*Planctomycetes-Verrucomicrobia-Chlamydiae*) bacteria. However, a decisive PVC-based eukaryogenesis scenario has not yet been reported. Here, we isolated PVC bacteria, unique for dividing by budding and for possessing developed endomembrane systems, from the deep sea. In cultured PVC bacterial strains, we detected typical eukaryotic organelle-like structures including endoplasmic reticulum, Golgi apparatus, vesicles, vacuoles and actin/tubulin-based microfilaments. Strikingly, we observed a nucleolus-containing nucleus in a *Verrucomicrobia* strain, which divides by mitosis. Transcriptomic results further demonstrated abundant presence of genes associated with eukaryotic cellular processes including membrane fusion. We propose that the prominent capability of membrane fusion drives eukaryogenesis by enabling PVC bacteria to evolve eukaryotic cellular features.

## Introduction

Life on earth originated from two kinds of organisms: prokaryotes and eukaryotes^1^. Although eukaryotes are generally considered to have evolved from prokaryotes, how this occurred remains one of the greatest enigmas in biology^2-4^. At the time of eukaryogenesis, the last eukaryotic common ancestor (LECA) had acquired a nucleus enclosed by a double membrane, membranous intra-cellular structures, mitochondria, and actin/tubulin-based cytoskeletal structures^2^. Moreover, the LECA also underwent intricate biological processes such as apoptosis, intron splicing, and meiosis^2^. Two competing hypotheses seek to explain the origin of eukaryogenesis: the endosymbiotic and the autogenic^5,6^. According to the autogenesis theory, the LECA evolved from a prokaryote through a gradual increase in cellular complexity that slowly accumulated over time^1,2,7^, and development of an endomembrane system that drove the compartmentalization of the nucleus^6^. Conversely, the endosymbiotic hypotheses propose that a moderately large and anaerobic prokaryote engulfed aerobic bacteria, some of which escaped digestion and instead became stabilized as endosymbionts, eventually evolving into mitochondria^8,9^. Testing these theories has been difficult given the reliance of canonical models unlikely to capture the wider spectrum of prokaryote cell biology and the paucity of known intermediate stages in the prokaryote-to-eukaryote transition^1,10^. However, if eukaryotes evolved from prokaryotes, then there must have been viable organisms with intermediate cellular structures and life styles between prokaryotes and eukaryotes^1,10^, and investigating such intermediates therefore holds the potential to decode the mysterious eukaryogenesis process.

Members within the PVC superphylum (which also includes bacteria from *Lentisphaerae* and *Kirimatiellaeota* as well as some uncultured candidate phyla) are examples of bacteria with features usually associated with eukaryotes or archaea, or both^10-12^, and hence represent exceptions to how we define bacteria. Consistently, earlier work proposed that the ancestor of the PVC bacteria diverged by developing distinct features that would later become known as archaeal, eukaryotic, or shared between them^10^. The PVC bacteria are thought to be the sister taxa to the LAECA lineage and the root of the Archaea and eukaryotes^10^. However, the objectivity and accuracy of this scenario need to be further clarified through culturing PVC bacteria that possessing more significant eukaryotic signatures (e.g. nucleus and other complex membranous cellular organelles).

Given that the deep sea is one of the most likely environments to harbor intermediates between prokaryotes and eukaryotes^1^ and has been proposed to be the proper place of life origin and stepwise evolution^13^, we here set out to culture deep-sea PVC bacterial strains. We found that these presented with many typical eukaryotic organelle-like structures, and specifically, detected a nucleolus-containing nucleus in a *Verrucomicrobia* strain, which divides through a mitosis-like manner with the involvement of the nucleolus. Lastly, we discussed the possible pathways of PVC-based eukaryogenesis driven by cell fusion.

## Results and Discussion

### Abundant eukaryotic signatures observed in deep-sea PVC bacteria

To closely assess presence of eukaryotic cellular features, we cultured four PVC bacteria, namely two *Planctomycetes* strains, one *Verrucomicrobia* strain and one *Lentisphaerae* strain isolated from deep-sea cold seeps (Fig. S1A). Through TEM analyses, we concluded that *Planctomycetes* and *Verrucomicrobia* strains likely divide by budding (Fig. S1B) and that membrane structures in PVC bacteria are both more complex and extended than those of typical Gram-positive (Expanded Data Fig. S1C, panel 1) or -negative bacterium (Expanded Data Fig. S1C, panel 2). PVC bacteria also have a condensed and intact nucleoid-like structure (Expanded Data Fig. S1C, panels 3-5), as exemplified by a typical nucleus-like organelle containing a well-shaped nucleolus in the *Verrucomicrobia* strain ZRK36 (Expanded Data Fig. S1C, panel 6), similar to those in fungal and human cells (Expanded Data Fig. S1C, panels 7-8). Additionally, eukaryotic signatures including endoplasmic reticula (ER) like-, vesicle like-, or vacuole like-organelles are present in *Planctomycetes* strain ZRK32 (Expanded Data Fig. S1C, panel 4). Although the presence of the nucleus and other membrane-bound organelles are hallmark features of eukaryotic cells, a recent cross-species proteomic analysis of the prokaryote nuclear matrix structure identified a significant fraction of such proteins evolutionary related to counterparts in eukaryotes^14^. In line with emerging studies highlighting unique properties of PCV bacteria^12^, we conclude that PVC strains are differentiated from how bacteria are conventionally defined by their abundant eukaryotic signatures as well as a high structural complexity. Intrigued by the distinct cell biology we had detected in *Planctomycetes* strain ZRK32 and *Verrucomicrobia* strain ZRK36, we decided to focus on these two strains for further studies.

### Eukaryotic signatures of *Plancomycetes*

To explore its additional eukaryotic features, we cultured *Planctomycetes* strain ZRK32 in both basal and rich media and analyzed the cellar ultra-structures in ultrathin TEM sections. We clearly detected several eukaryotic membrane-trafficking machinery components including the Golgi apparatus and the ER (Fig. 1A and Expanded Data Figs. S2-S3). In the prokaryote-to-eukaryote evolutionary transition, the emergence of an endomembrane system resembling the Golgi and ER was a pivotal event^9^, as internal tubular networks enable specialization and compartmentalization of biochemical processes, as well as a varied surface matrix supporting rich intracellular processes and signaling cascades^15,16^. In eukaryotic cells, highly specialized vesicle trafficking protein complexes direct transport between the ER and Golgi^2^. In the strain ZRK32, we identified both a vesicle like-structure (Fig. 1A and Expanded Data Figs. S4A-C) and a vacuole-like structure (Fig. 1A and Expanded Data Figs. S4D-F). Vacuoles are highly dynamic and pleiomorphic^17^, with a size that varies depending on the cell type and growth conditions (Expanded Data Figs. S4D-F). Vacuoles store cellular components such as proteins, sugars etc., and play essential roles in plants response to different biotic/abiotic signaling pathways^17^. The presence of vacuoles in strain ZRK32 suggests that *Planctomycetes* bacteria might have adopted a eukaryotic mechanism for nutrient metabolism and signal transduction. In the TEM sections, we also observed several types of actin/tubulin-derived fibrous structures (Fig. 1A, panels 7 and 8), suggesting that *Planctomycetes* members also evolved sophisticated cytoskeletal machineries, such as those present in eukaryotes. These are also present in other *Planctomycetes* strains, and proposed to be involved in specific locomotion and engulfment processes^18^.

**Fig. 1.**
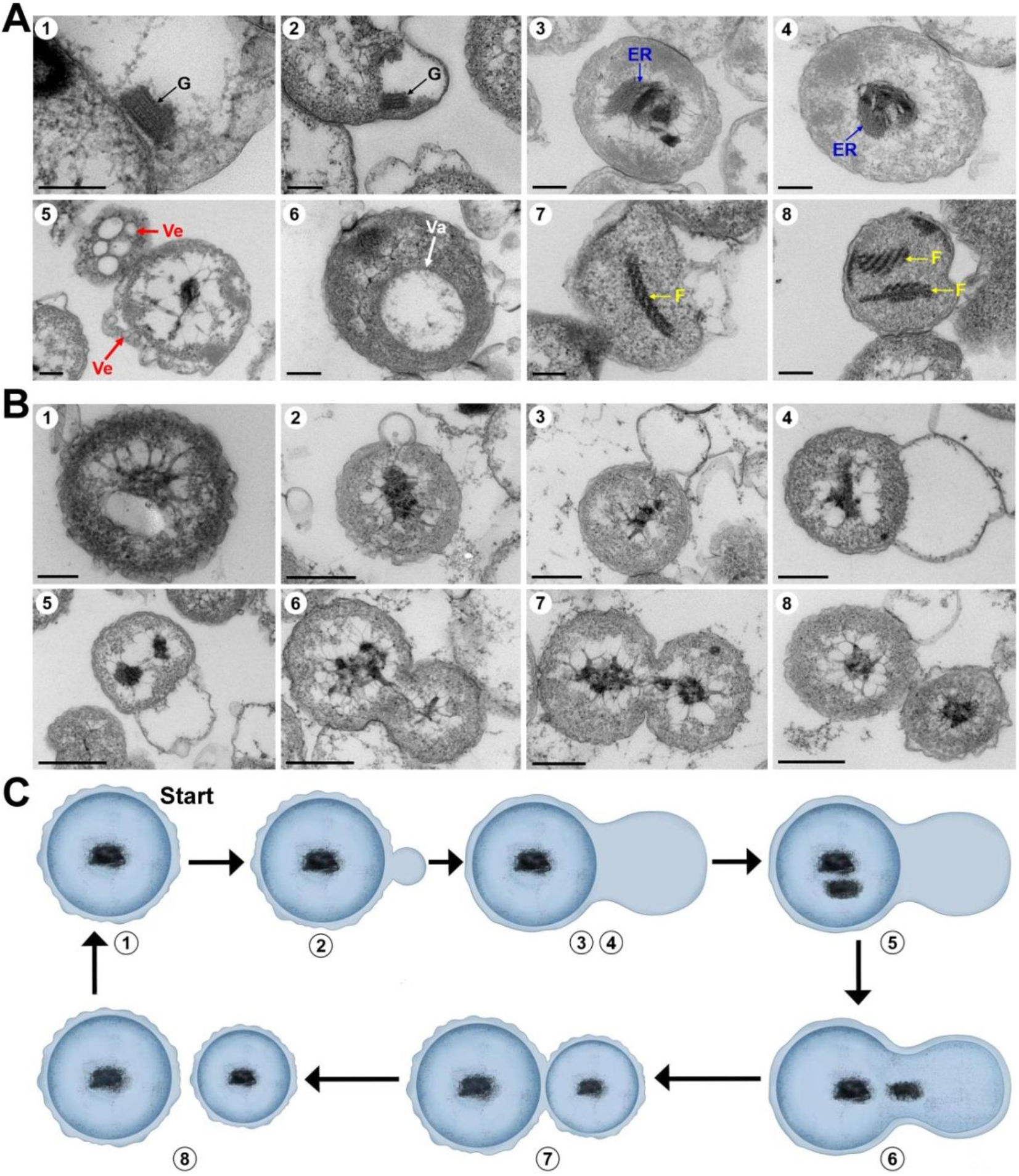
The unique eukaryotic signature and cell division pattern of *Planctomycetes* strain ZRK32. (**A**) Ultrathin TEM sections showing Golgi apparatus like- (indicated with G in panels 1 and 2), endoplasmic reticula like- (indicated with ER in panels 3 and 4), vesicle like- (indicated with Ve in panel 5), vacuole like- (indicated with Va in panel 6), and microfilaments-like (indicated with F in panels 7 and 8) organelles observed in the cell of *Planctomycetes* strain ZRK32. (**B**) Ultrathin TEM sections showing the process of particular polar budding division (panels 1-8) conducted by strain ZRK32. Representative pictures representing different phases of cell division are shown. Scale bars are 200 nm in panels A and B. (**C**) The proposed model of cell division of *Planctomycetes* strain ZRK32 based on the TEM observation shown in panel B. The numbers are the same to those shown in panel B.

The *Planctomycetes* strain ZRK32 is shown to divide by budding as previously reported^19^. In the beginning of division, the membrane extends and forms a bulge region, which becomes larger until its size is similar to that of the mother cell (Figs. 1B and 1C, panels 1 to 4). Meanwhile, the genetic materials within the nucleoid are duplicated and equally divided between the mother and daughter cells along with other cytoplasmic contents (Figs. 1B and 1C, panels 5 to 8), after which the daughter cell completely separates from the mother cell to finalize a cell division cycle. Here, we asked whether genes associated with eukaryotic features and cell division are functional during growth. To this end, we performed a comparative transcriptomic analysis of strain ZRK32 cultured in basal and rich media (Fig. 2A). Overall, the transcriptomic profiles revealed significant regulation of genes encoding ER associated proteins, Golgi vesicle transport, vacuole sorting, SNARE-associated Golgi protein, cadherin domain, and Rab/Rho GTPases (Fig. 2B). While SNARE-associated proteins play specific roles in vesicle fusion^20^, the small Ras GTPase superfamily, one of the largest protein families in eukaryotes, is broadly involved in various regulatory processes, including cytoskeleton remodelling, signal transduction, nucleocytoplasmic transport and vesicular trafficking^21^. Additionally, genes associated with actin/tubulin are strongly upregulated (Fig. 2C). In eukaryotes, both actin and tubulin polymerize to form filaments that plays central roles in cellular motility, cytoskeleton formation and phagocytosis^22^. Their presence and upregulation in strain ZRK32 indicate that filament structures might play a critical role in determining its cell shape and movement. In microbial cell division, members of the tubulin-homologue Fts protein family assemble at the future site of cell division to form a contractile ring, known as the Z ring^23^. Consistent with earlier work^19^, we did not detect the FtsZ isoform, but numerous other Fts-proteins were present and upregulated in rich medium-supported growth (Fig. 2D). Within the group of proteins associated with cytoskeletal function, the gene encoding for VgrG1, an actin cross-linking protein known for a role in the bacterial type VI secretion system, to be the most upregulated^24^. We also found that the bacterial actin-analogue MreB, was upregulated in strain ZRK32 (Fig. 2C). Peptidoglycan ultimately determine bacterial cell shape, but shape is also influenced by cytoskeletal filaments such as MreB that participate in its formation and degradation^25^. MreB is probably not generally required for either cell division or cell-shape determination in *Planctomycetes*^19^, but its function in cellular processes is diversified in different *Planctomycetes* strains.

**Fig. 2.**
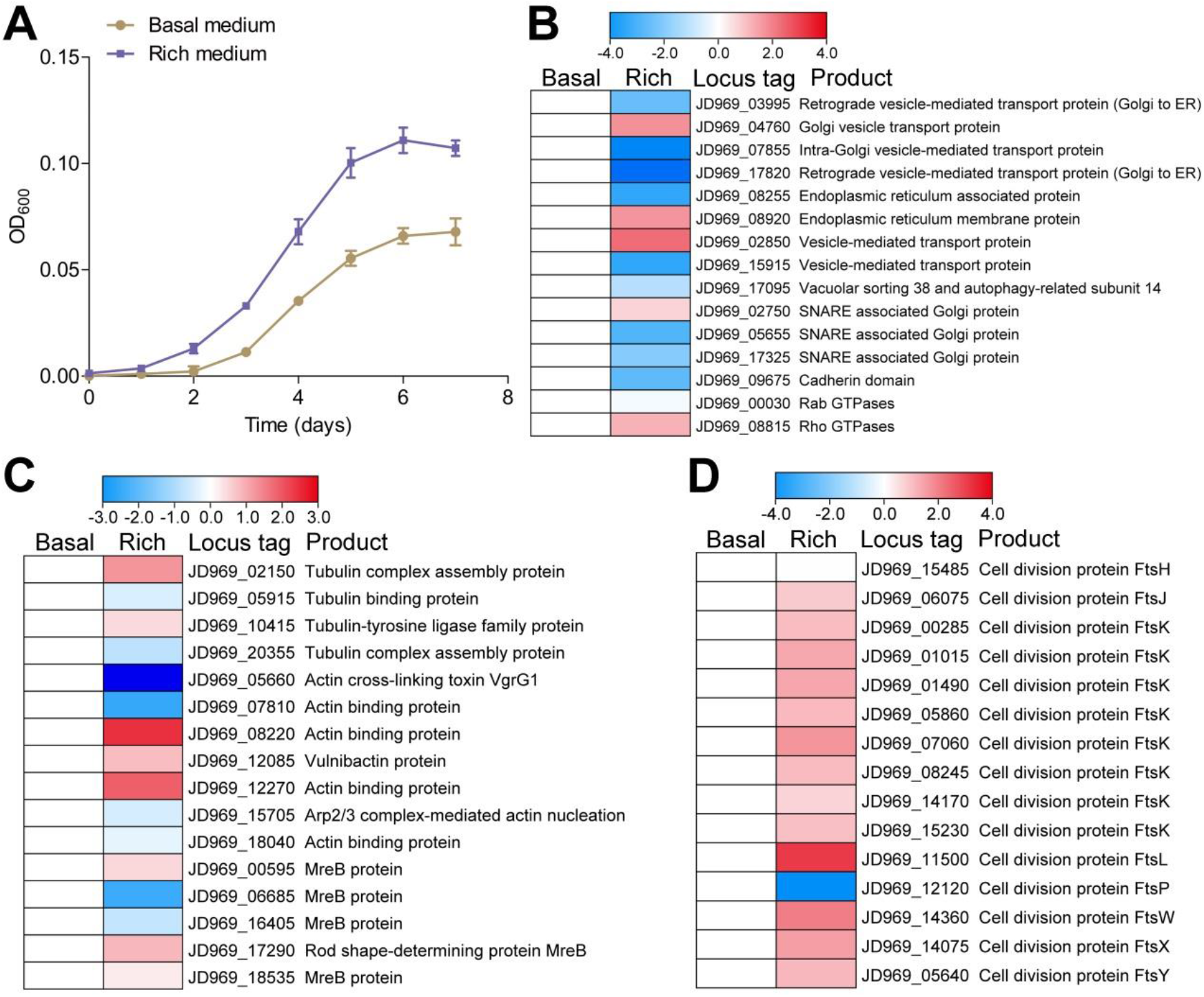
Transcription assays of genes associated with eukaryotic signatures and cell division of *Planctomycetes* strain ZRK32. (**A**) Growth assays of strain ZRK32 cultivated in either rich medium or basal medium. (**B**) Transcriptomics based heat map showing differentially expressed genes encoding proteins associated with eukaryotic signatures in strain ZRK32. (**C**) Transcriptomics based heat map showing differentially expressed genes encoding proteins associated with cytoskeleton in strain ZRK32. (**D**) Transcriptomics based heat map showing differentially expressed genes encoding key proteins associated with cell division in strain ZRK32.

### Eukaryotic signatures of *Verrucomicrobia*

Similar to what we observed in *Planctomycetes* strain ZRK32, ultrathin TEM sections of *Verrucomicrobia* strain ZRK36 cells revealed presence of ER like-and vacuole like-structures (Fig. 3A). However, the most striking feature of *Verrucomicrobia* strain ZRK36 is its well-shaped nucleus containing a nucleolus-like structure (Fig. 3A, panels 1-2). Although the nucleoid of *Planctomycetes* is considered to be more like eukaryotic nuclei^26^, the kind of clear and intact nucleus structure that we observed in strain ZRK36 has to our knowledge not been reported in any other bacteria. The nucleus, which contains almost all of the cells’ genetic material, is a remarkably complex structure, and a hallmark feature of eukaryote cells^26,27^. Strikingly, we also observed a structure resembling the nucleolus (Fig. 3A, panels 1-3), whose main function in eukaryote cells is ribosome biogenesis and ribosomal RNA (rRNA) synthesis^28^. In the initial stage of budding division, the nucleolus in strain ZRK36 is duplicated and the new copy along with cytoplasmic materials enters into the newly formed budding structure (Fig. 3B, panel 2). Thereafter, the size of budding becomes almost equal to that of the mother cell (Fig. 3B, panel 3). At that stage, the genetic materials inside the nucleolus are disassembled, thereby directing development of the daughter cell (Fig. 3B, panels 4 and 5). Thereafter, the formation of a new nucleus is initiated, as demarcated by a condensed nucleolus (Fig. 3B, panel 6). Subsequently, the spindle fibers between the old and new nuclei are extended along with separation of the two nuclei (Fig. 3B, panel 7). Finally, the extended spindle fibers disappear and the daughter cell becomes thoroughly segregated from the mother cell, completing the division process (Fig. 3B, panels 8-10). At the end of mitosis, eukaryotic cells reshape the intact nucleus by segregating the copies of the replicated genome into two new nuclear compartments, which splits into two in a “closed mitosis”^29^. Here, we found that the ZRK36 strain undergoes a meiosis-like cell division process that is highly similar to that of budding yeast (Figs. 3C and 3D), supporting the conclusion that this PVC bacterium evolved to adapt a eukaryotic cell division mode. Meiosis is widely considered to be a core process unique to eukaryotes, and consequently expected to be absent from all known prokaryote^2^. However, on the basis of our high-resolution morphology analysis of strain ZRK36 in ultrathin TEM sections, we conclude that some PVC bacteria throughout evolution acquired eukaryotic organelle-like structures as well as cell cycle progression and cell division modes believed to be solely conducted in eukaryotes.

**Fig. 3.**
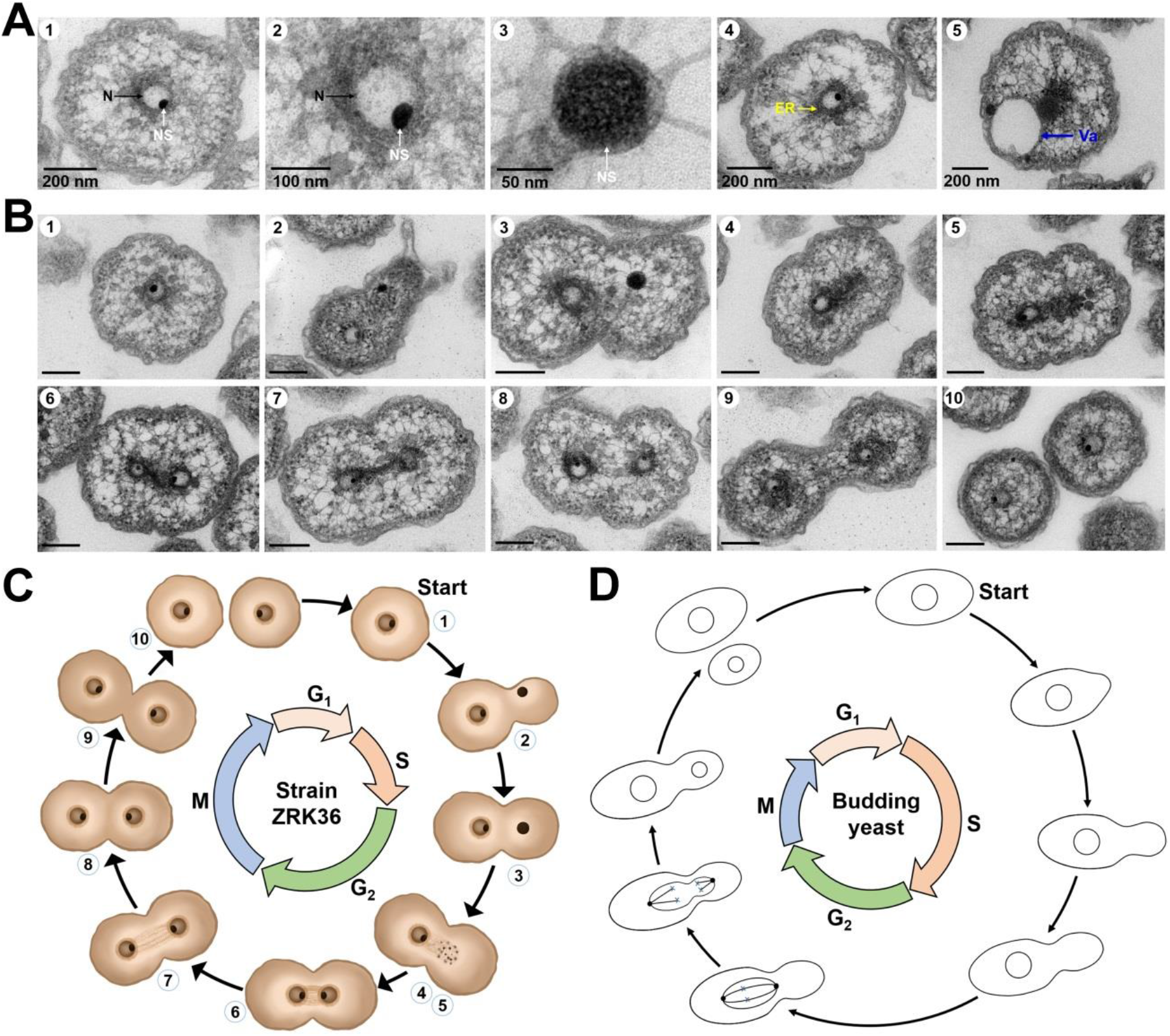
The unique eukaryotic signatures and cell division pattern of *Verrucomicrobia* strain ZRK36. (**A**) Ultrathin TEM sections showing, nucleus like- (indicated with N in panels 1 and 2), nucleolus like- (indicated with NS in panels 1 to 3), endoplasmic reticula like- (indicated with ER in panel 4) and vacuole like- (indicated with Va in panel 5) organelles observed in the cell of *Verrucomicrobia* strain ZRK36. (**B**) Ultrathin TEM sections showing the process of a typical mitosis manner (panels 1-10) conducted by *Verrucomicrobia* strain ZRK36. Representative pictures representing different phases of cell division are shown. Scale bars are 200 nm. (**C**) The proposed model of cell division conducted by *Verrucomicrobia* strain ZRK36 based on the TEM observation shown in panel B. The numbers are the same to those shown in panel B. (**D**) Typical cell division process conducted by the budding yeast. G1, S, G2 and M in panels C and D represent different phases of meiosis.

It’s noteworthy that many genes that encode factors associated with eukaryotic organelles and unique cell biology are significantly regulated during growth (Fig. 4B), strongly suggesting that they function similarly in processes that contribute to the growth of strain ZRK36 (Fig. 4A). Among growth-regulated genes, *NOG1*, which encodes the nucleolar GTPase Nog1, a coordinator factor in ribosome formation, is known to be present in the nucleolus in humans and yeast^30,31^. In yeast, the *NOG1* gene is essential for cell viability and plays an important role in nucleolar functions^30^. Based on its described function in eukaryotic cells, we propose that the NOG1 homologue in strain ZRK36 plays a similar nucleolus-directed role in cell division. Similarly, we found that the gene encoding centromere protein H (CENPH) is also regulated, implicating mitosis as another distinct eukaryotic process in strain ZRK36 (Fig. 4B). CENPH is as a component of the active centromere complex, which directs kinetochore assembly on the centromeric chromatin to ensure faithful chromosome segregation during mitosis^32,33^. Given the combined electron microscopy analysis and transcriptomic profiling, we propose that strain ZRK36 undergoes a eukaryotic-like mitosis process.

**Fig. 4.**
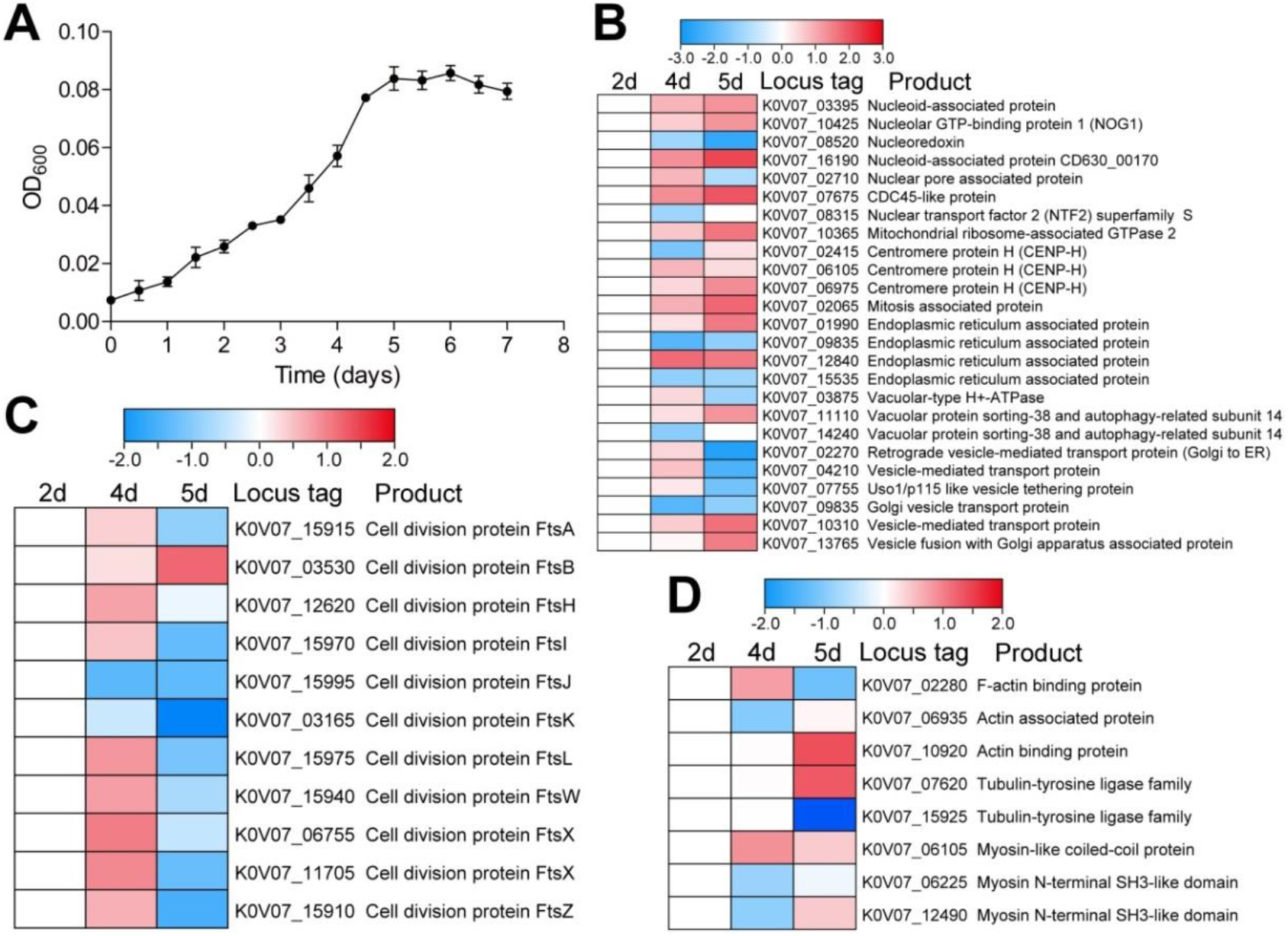
Transcription assays of genes associated with eukaryotic signatures and cell division of *Verrucomicrobia* strain ZRK36. (**A**) Growth assays of strain ZRK36 cultivated in basal medium. **(B)** Transcriptomics based heat map showing differentially expressed genes encoding proteins associated with eukaryotic signatures in strain ZRK36. (**C**) Transcriptomics based heat map showing differentially expressed genes encoding key proteins associated with cell division in strain ZRK32. (**D**) Transcriptomics based heat map showing differentially expressed genes encoding proteins associated with cytoskeleton in strain ZRK36.

In contrast with *Planctomycetes* strain ZRK32, FtsZ is present in *Verrucomicrobia* ZRK36 where it likely participates in the process of cell division together with other Fts-proteins (Fig. 4C) and actin/tubulin associated proteins (Fig. 4D). Since FtsZ is a tubulin family protein, it is conceivable that early gene duplication giving rise to both tubulins and FtsZ occurred in some *Verrucomicrobia* bacteria. Taken together, our observations of abundant eukaryotic features in strains ZRK32 and ZRK36 illustrate how deep-sea PVC bacteria challenge the knowledge we have gained by studying conventional model organisms and suggest that these provide good models to explore eukaryogenesis.

### Significant membrane fusion capability of PVC bacteria

A feature characteristic of PVC bacteria, but not typically seen in other bacteria, is their endomembrane system, which is particularly well developed in some species^12,23,34^. Unlike in prokaryotes, fusion of membranous organelles is the basis of life in eukaryotes, with numerous processes hinging on the flawless timing and operation of membrane fusion^35^. Notably, all deep-sea PVC members cultured in this study have well-developed endomembrane system and show diverse abilities of membrane fusion. In the *Lentisphaerae* strain zth2, we captured several extracellular protrusions that mediate cell-cell interactions and membrane fusion (Fig. 5A), which enables the formation of an integrated channel for exchanging cellular material (Fig. 5A). Indeed, in the inside-out theory of eukaryote evolution, protrusions that enable material exchange are suggested to function as a cellular mechanism that engulfs mitochondria-like bacteria^6^. Here, we find that *Planctomycetes* strain ZRK32 has a remarkable endocytosis capability, enabling vesicle formation from the extracellular environments via cell fusion and forming a larger size vesicle (Fig. 5B). We speculate that superiority in capturing extracellular nutrient and phagocytose smaller cells might endow endomembrane system-carrying bacteria with both an evolutionary and survival advantage. In *Verrucomicrobia* strain ZRK36, we detect a dramatic cell fusion capability, allowing the formation of giant cells with several-fold size of the original cell (Expanded Data Fig. 5C and Fig. S5). In addition, several nuclei-like organelles are capable of fusing into a bigger size nucleus (Fig. 5C, panel 6). Moreover, giant cells are more prevalent in the stationary phase than in the logarithmic phase.

**Fig. 5.**
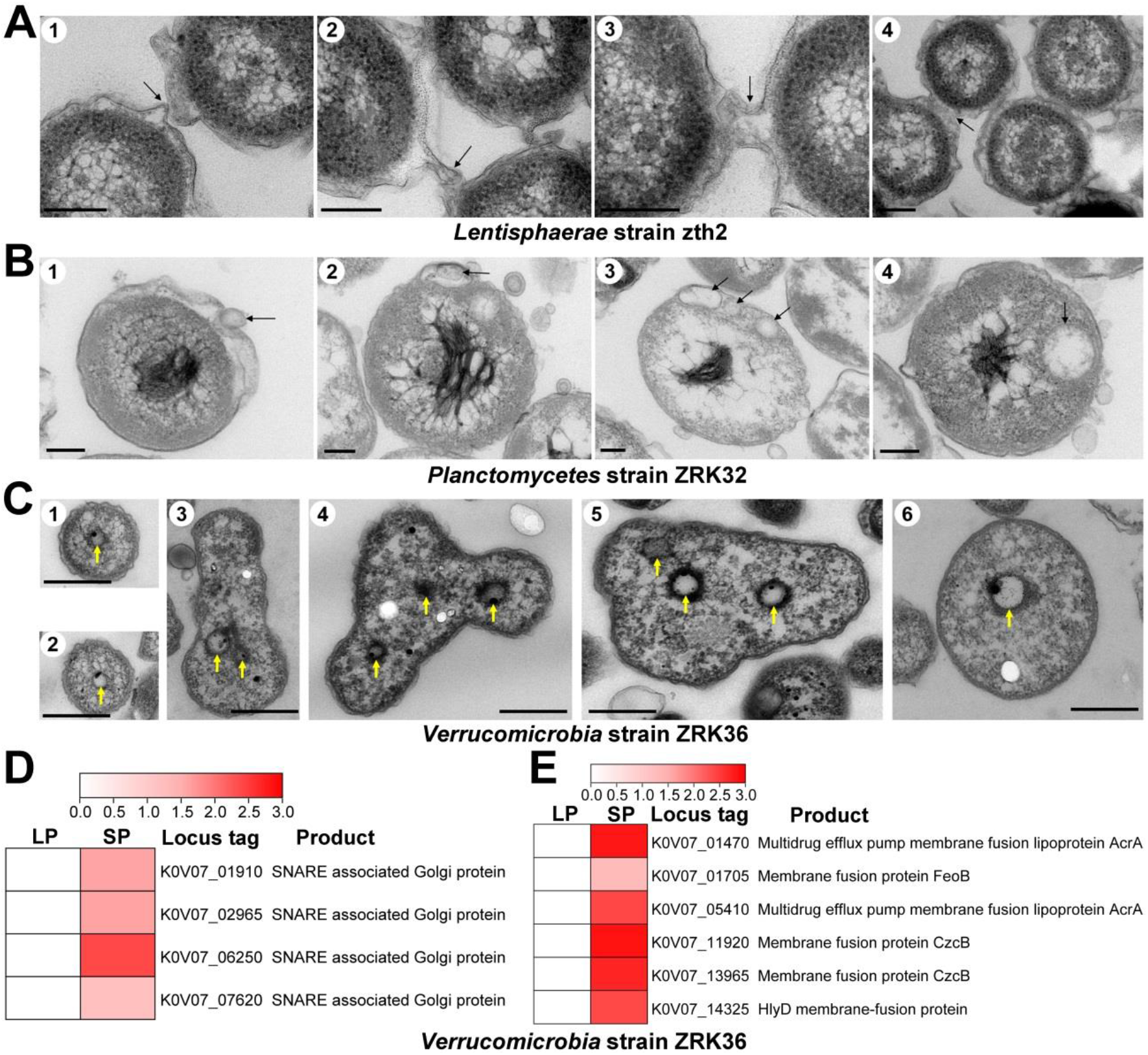
Diverse membrane fusion manners conducted by PVC bacteria. (**A**) Ultrathin TEM sections showing the process of membrane fusion directed by the protrusions around the cell of *Lentisphaerae* strain zth2. The arrows in panels 1 to 4 indicate the position of membrane fusion. (**B**) Ultrathin TEM sections showing the endocytic process through a membrane fusion manner conducted by *Planctomycetes* strain ZRK32. The arrows in panels 1 to 4 indicate the positions of vesicle. (**C**) Ultrathin TEM sections showing the inter-species phagocytosis conducted by *Verrucomicrobia* strain ZRK36. The arrows in panels 1 to 6 indicate the positions of nucleus like-organelles. (**D**) Transcriptomics based heat map showing differentially expressed genes encoding SNARE-associated Golgi proteins in *Verrucomicrobia* strain ZRK36. (**E**) Transcriptomics based heat map showing differentially expressed genes encoding proteins associated with membrane fusion in *Verrucomicrobia* strain ZRK36. “LP” indicates the logarithmic phase; “SP” indicates the stationary phase.

Consistent with our phenotypic analysis (Fig. 5C), the transcriptome profiles reveal that membrane fusion genes are significantly upregulated (Figs. 5D and 5E). Among these, four genes encoding SNARE-associated proteins are prominently up-regulated in the stationary phase relative to the logarithmic phase (Fig. 5D). Membrane fusion, a universal feature of eukaryotic protein trafficking, is mediated by SNARE-associated proteins that embed in opposing membranes to spontaneously drive membrane fusion and cargo exchange^36^. Additionally, the expressions of other membrane fusion proteins/lipoproteins were also up-regulated in the stationary phase (Fig. 5E). Based on these analyses, we conclude that *Verrucomicrobia* strain ZRK36 has evolved a significant cell fusion capability, enabling it to form more complicated and larger cells, and executing processes resembling phagocytosis, an eukaryotic innovation conserved from unicellular protists to animals, that enabled eukaryotes to feed on other organisms^18^. Presumably, other PVC bacteria might also evolve similar membrane fusion strategies to increase cell complex and size, and in turn facilitating eukaryogenesis.

### PVC-based eukaryogenesis

PVC bacteria have long been deemed close associates of the eukaryote ancestor^12,23,26^. Although eukaryogenesis-predictive features are common in PVC bacteria, their implications and real functions have not been adequately addressed. Based on our data and recent related work, we speculate that eukaryogenesis could have evolved in PVC bacteria through two core mechanisms: inter-species and cross-species fusions. In inter-species fusion, two PVC bacteria, each with a well-developed nucleus (e.g. *Verrucomicrobia* strain ZRK36) first fuses to form a larger cell, which then sequentially fuses with bacteria that has acquired either eukaryote-like endomembrane systems or other organelles to form a complex eukaryotic cell (Fig. 6A). In cross-species fusion, a PVC bacterium with a developed nucleus first fuses with a PVC or a related strain to form a larger cell with developed eukaryotic features, which then sequentially fuses with bacteria that has acquired a nucleus-like or other organelle-like structures to form a complex eukaryotic cell (Fig. 6B). In the hypothetical LECA, the endomembrane systems, including the ER, Golgi complex and the main components of the endocytosis and recycling systems, were already present ^2^, but the evolutionary events preceding this transition are a matter of debate. Based on comparative genomic analysis, earlier work proposed the Asgard superphylum, belonging to archaea (a separate domain from bacteria and eukaryotes that lack nuclei), as a candidate host cell that merged with a bacterial mitochondrial endosymbiont to generate the first eukaryotic cell^37^. This scenario supports a 2D model of evolution in which bacteria and archaea evolved along two separate branches of the tree of life, and eukaryotes evolved from archaea^37^. Here, at a cell biological and transcriptome level, we discovered more developed eukaryotic features in deep-sea PVC bacteria than previously reported, most notably a nucleolus-containing nucleus and mitosis, supporting the alternative and recent hypothesis of PVC-based eukaryogenesis^10^. Based on our data, we propose a modified PVC-based linear (1D) scenario of eukaryogenesis (Fig. 6C). In our model, we propose that the last universal common ancestor (LUCA) diverged into two linages: bacteria with very few archaeal and eukaryotic features, and the last PVC common ancestor (LPCA), with abundant archaeal or eukaryotic features, or both. At the subsequent evolutionary branch point, the LPCA diverged into the last modern PVC common ancestor (LMPCA) that had lost most eukaryotic signatures, and the last archaeal and eukaryotic common ancestor (LAECA). In this scenario, PVC bacteria are related to intermediates in both LPCA derived lineages: LMPCA and LAECA, the former less complex, and the latter more complex, progressively accumulating eukaryotic or archaeal features and thereby increasing its cellular complexity. However, intermediates diverging towards LMPCA or LEACA likely share many common features initially, resulting in universal communication between these two groups through cell fusions (as indicated with dotted lines between the LMPCA and LEACA branches). Indeed, findings in recent years revealed that eukaryotic ancestors more easily exchanged genes when they were more closely related, particularly at the start of their diversification^2^. Throughout PVC-based eukaryogenesis, ubiquitous and multilateral fusions would be expected to create chimeric cells containing increasingly complicated genetic materials and other organelle-like structures. We propose that the PVC members cultured in the present study (*Planctomycetes* strain ZRK32 and *Verrucomicrobia* strain ZRK36) might represent intermediate states between the LPCA and the LAECA as the LPCA became more “eukaryotic” with a gain of complexity (Fig. 6C)^10^. The LPCA as well as intermediates between the LPCA and the LAECA will unlikely have evolved a sophisticated nucleus or eukaryote-associated features^38^; instead these stages are likely evolutionary “works in progress” until the very end of eukaryogenesis, which remains as the main gap in the history of life on the Earth (Fig. 6C). Hence, we propose that future exploration of prokaryotic diversity should focus on the later time line between the LPCA and the LAECA, which diverged to the last archaeal common ancestor (LACA) and the last eukaryotic common ancestor (LECA). In the PVC-based linear tree of life, archaea have lost most of the eukaryotic features, with Asgard having lost the least and diverged less^10^, consistent with their sheer number of eukaryotic features and the phylogenetic proximity of these eukaryotic features to eukaryotes (Fig. 6C)^37^. The PVC-based 1D scenario we outlined here does not deny the proximity of Asgard to Eukarya, nor does it question the notion that eukaryotes may have evolved from an archaeal ancestor. Instead, the PVC-based linear scenario addresses most of the inconsistencies of current 2D scenarios, and properly integrates most recent discoveries, providing a coherent path of evolution of the three domains of life.

**Fig. 6.**
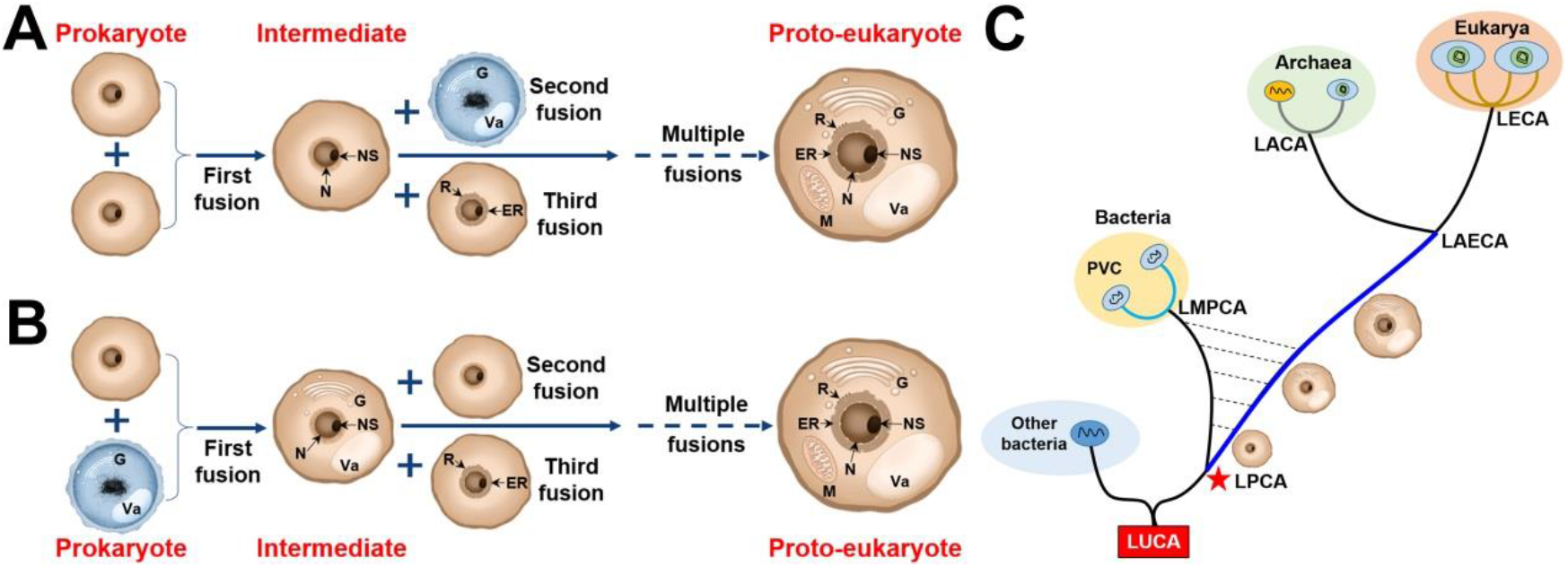
Cell fusion drives PVC-based eukaryogenesis. (**A**) A proposed model of inter-species fusion of PVC bacteria launching the stepwise eukaryogenesis. ER, endoplasmic reticula; N, nucleus; NS, nucleolus; G, Golgi apparatus; Va, vacuole; M, mitochondria; R, ribosome. (**B**) A proposed model of cross-species fusion of PVC bacteria launching the stepwise eukaryogenesis. (**C**) PVC-based 1D tree of life. LUCA, the last universal common ancestor; LPCA, the last PVC common ancestor; LMPCA, the last modern PVC common ancestor; LAECA, the last archaeal and eukaryotic common ancestor; LACA, the last archaeal common ancestor; LECA, the last eukaryotic common ancestor.

## Methods

### Isolation and cultivation of deep-sea PVC bacteria

The PVC bacterial strains (including TYQ1, ZRK32, ZRK36 and zth2) were isolated from deep-sea deposits obtained from a typical cold seep in the South China Sea by *RV KEXUE*^*39,40*^. Strain TYQ1 was obtained from an inoculum enriched in the aerobic MF7 medium (containing 2.38 g/L HEPES as a buffer, 5 mL/L vitamin solution, 1 mL/L trace element solution, 0.25 g/L glucose, 0.25 g/L peptone, 0.25 g/L yeast extract, 1 g/L N-acetylglucosamine, 200 mg/L ampicillin, 100 mg/L carbenicilli, 200 mg/L streptomycin, 20 mg/L amphotericin in 1 L filtered seawater, pH=7.0). The enrichment was incubated at 28 °C for one week. Thereafter, enrichments were plated onto the MF7 medium supplemented with 15 g/L agar. Then small colorless colonies were observed after one week and the strain TYQ1 was purified by continuous re-streaking on MF7 solid plates until it was considered to be pure. Strains ZRK32, ZRK36 and zth2 were obtained from enrichments incubated in the anaerobic basal medium (containing 1.0 g/L yeast extract, 1.0 g/L peptone, 1.0 g/L NH_4_Cl, 1.0 g/L NaHCO_3_, 1.0 g/L CH_3_COONa, 0.5 g/L KH_2_PO_4_, 0.2 g/L MgSO_4_.7H_2_O, 0.7 g/L cysteine hydrochloride, 500 µL/L 0.1 % (w/v) resazurin in filtered seawater, pH=7.0) supplemented with 50 mg/L rifampicin or 1 g/L laminarin at 28 °C for one month. Then 50 µL enrichment was spread on Hungate tubes covered by the basal medium supplemented with 15 g/L agar and incubated at 28 °C for 10 days. Individual colonies with distinct morphology were picked using sterilized bamboo sticks and then cultured in the basal medium supplemented with 50 mg/L rifampicin or 1 g/L laminarin by repeated use of the Hungate roll-tube methods for several rounds until they were considered to be pure^41,42^. The purity of all strains (TYQ1, ZRK32, ZRK36 and zth2) were confirmed by Transmission electron microscopy (TEM) and repeated partial sequencing of the 16S rRNA gene, then these pure bacterial strains were preserved at -80 °C with 20% (v/v) glycerol.

### Phylogenetic analysis

The full-length 16S rRNA gene sequences of four cultured PVC bacterial strains (TYQ1, ZRK32, ZRK36 and zth2) were obtained from their genomes and deposited in the GenBank database, and other bacteria used for phylogenetic analysis were obtained from NCBI (www.ncbi.nlm.nih.gov). All the sequences were aligned by MAFFT version7^43^ and manually corrected. The phylogenetic tree was constructed under the W-IQ-TREE web server (http://iqtree.cibiv.univie.ac.at)^44^ with GTR+F+I+G4 model. And then this tree was edited by the online tool Interactive Tree of Life (iTOL v5)^45^.

### TEM observation

To observe the morphological characteristics of PVC bacteria, the cell suspension was washed with the PBS buffer (containing 137 mM NaCl, 2.7 mM KCl, 10 mM Na_2_HPO_4_, 1.8 mM KH_2_PO_4_, 1 L sterile water, pH 7.4) and centrifuged at 5,000 × *g* for 10 min, and taken by immersing copper grids coated with a carbon film for 30 min, washed in the PBS buffer and air dried. For ultrastructure observation of PVC bacteria, ultrathin-section electron microscopy was performed as described previously^46^. Briefly, four PVC strains were grown to the stationary phase in the rich medium and then centrifuged at 5,000 × *g* for 10 min. The samples were firstly preserved in 2.5% (v/v) glutaraldehyde overnight at 4 °C, washed three times with the PBS buffer and dehydrated in ethanol solutions of 30%, 50%, 70%, 90% and 100% for 10 min each time, and then the samples were embedded in a plastic resin. Ultrathin sections (50∼70 nm) of cells were prepared with an ultramicrotome (Leica EM UC7, Wetzlar, Germany), stained with uranyl acetate and lead citrate. The ultrathin-sections of *Bacillus thuringiensis* A260, *Pseudomonas stutzeri* 273, *Magnaporthe grisea* and human liver cancer Huh 7.5 cell line were also prepared accordingly to the above procedures. All samples were observed using TEM (HT7700, Hitachi, Tokyo, Japan) with a JEOL JEM 12000 EX (equipped with a field emission gun) at 100 kV^39^.

### Genome sequencing and assembly

Whole-genome sequence determinations of four PVC strains (TYQ1, ZRK32, ZRK36 and zth2) were carried out with the Oxford Nanopore MinION (Oxford, UK) and Illumina MiSeq sequencing platform (San Diego, California, USA). A complete description of the sequencing assembly and subsequent analyses was performed as previously described^39^.

### Growth assay and transcriptomic analysis of strain ZRK32

To assess the effect of organic nutrition on the growth of strain ZRK32, 15 mL freshly cells were inoculated in 1.5 L basal medium and rich medium at 28 °C for 7 days, respectively. Each condition was performed three times. Bacterial growth status was monitored by measuring the OD_600_ value each day until the cell growth reached the stationary phase. For transcriptomic analysis, cell suspensions of strain ZRK32 cultured in 1.5 L basal medium or rich medium for 6 days were collected at 8,000 × *g* for 20 min. The transcriptomic sequencing was performed by Novogene (Tianjin, China), and detailed protocols including library preparation, clustering and sequencing and data analyses were performed as previously described^39^.

### Growth assay and transcriptomic analysis of strain ZRK36

Growth assays of strain ZRK36 were performed at atmospheric pressure. Briefly, 15 mL fresh culture of strain ZRK36 was inoculated in 2 L Hungate bottles containing 1.5 L basal medium, and then anaerobically incubated at 28 °C for 7 days. And bacterial growth status was monitored by measuring the OD_600_ value every 12 h until the cell growth reached the stationary phase. For transcriptomic analysis, cell suspensions of strain ZRK36 cultured in 1.5 L basal medium for 2, 4, 5 days were collected at 8,000 × *g* for 20 min, respectively. Then, the transcriptomic sequencing and analysis were performed by Novogene (Tianjin, China) as previously described^39^.

## Data deposits and availability

The raw amplicon sequencing data have been deposited to NCBI Short Read Archive (accession numbers: PRJNA768640). The full-length 16S rRNA gene sequences of PVC superphylum strains (TYQ1, ZRK32, ZRK36 and zth2) have been deposited at GenBank under the accession numbers: OK160720, MW376756, MZ646149 and MW729759, respectively. The complete genome sequences of PVC superphylum strains (TYQ1, ZRK32, ZRK36 and zth2) have been deposited at GenBank under the accession numbers: CP083740, CP066225, CP080649 and CP071032, respectively. The raw sequencing reads from the transcriptomics analysis of strain ZRK32 incubated in basal medium and rich medium have been deposited to the NCBI Short Read Archive (accession numbers: PRJNA768630). The raw sequencing reads from the transcriptomics analysis of strain ZRK36 at different stages of growth have been deposited to the NCBI Short Read Archive (accession numbers: PRJNA768634 and PRJNA769601).

## Acknowledgements

We thank Dr. Christina Lilliehook, from the professional editing group “Life Science Editors”, for improving the scientific and writing quality of this manuscript. This work was funded by the Strategic Priority Research Program of the Chinese Academy of Sciences (Grant No. XDA22050301), the Major Research Plan of the National Natural Science Foundation (Grant No. 92051107), Key deployment projects of Center of Ocean Mega-Science of the Chinese Academy of Sciences (Grant No. COMS2020Q04), National Key R&D Program of China (Grant No. 2018YFC0310800) and the Taishan Young Scholar Program of Shandong Province (tsqn20161051) for Chaomin Sun.

## Author contributions

RZ and CS conceived and designed the study; RZ conducted most of the experiments; CW helped to do some experiments related to microscopic observation; TZ helped to isolate the *Lentisphaerae* bacterium; YT helped to isolate the *Planctomycetes* bacterium; RZ and CS lead the writing of the manuscript; all authors contributed to and reviewed the manuscript.

## Expanded Data

**Expanded Data Fig. S1.**
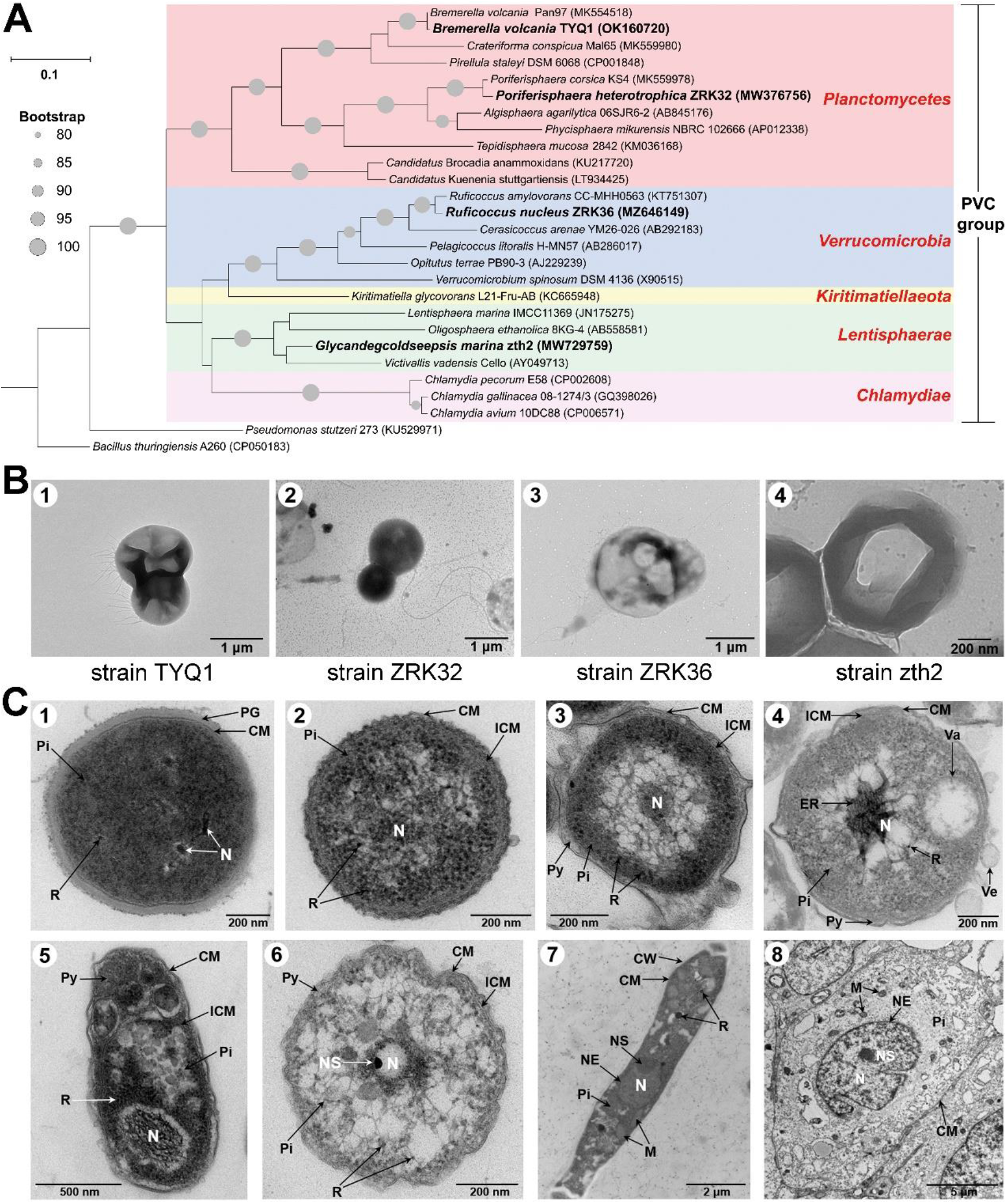
Phylogeny and morphology of deep-sea novel members belonging to the PVC superphylum. **(A)** 16S rRNA-based maximum-likelihood phylogenetic tree shows the positions of novel deep-sea *Lentisphaerae, Planctomycetes* and *Verrucomicrobia* strains are clustered within the PVC group. Bootstrap values (%) > 80 are indicated at the base of each node with the gray dots (expressed as percentages of 1,000 replications). The access number of each 16S rRNA is indicated after the strain’s name. The sequences of *Pseudomonas stutzeri* 273 and *Bacillus thuringiensis* A260 are used as outgroup. Bar, 0.1 substitutions per nucleotide position. **(B)** TEM observation of morphology of deep-sea *Planctomycetes* strains TYQ1 (panel 1) and ZRK32 (panel 2), *Verrucomicrobia* strain ZRK36 (panel 3) and *Lentisphaerae* strain zth2 (panel 4).**(C)** Ultrathin TEM sections of *Bacillus thuringiensis* A260 (panel 1), *Pseudomonas stutzeri* 273 (panel 2), *Lentisphaerae* strain zth2 (panel 3), *Planctomycetes* strains ZRK32 (panel 4) and TYQ1 (panel 5), *Verrucomicrobia* strain ZRK36 (panel 6), *Magnaporthe grisea* (panel 7) and human liver cancer Huh 7.5 cell line (panel 8). PG, peptidoglycan; CM, cytoplasmic membrane; Pi, pirellulosome; R, ribosome; N, nucleoid or nucleus; ICM, intracytoplasmic membrane; Py, paryphoplasm; Va, vacuole like-organelles; NS, nucleolus; NE, nucleus envelope; M, mitochondria.

**Expanded Data Fig. S2.**
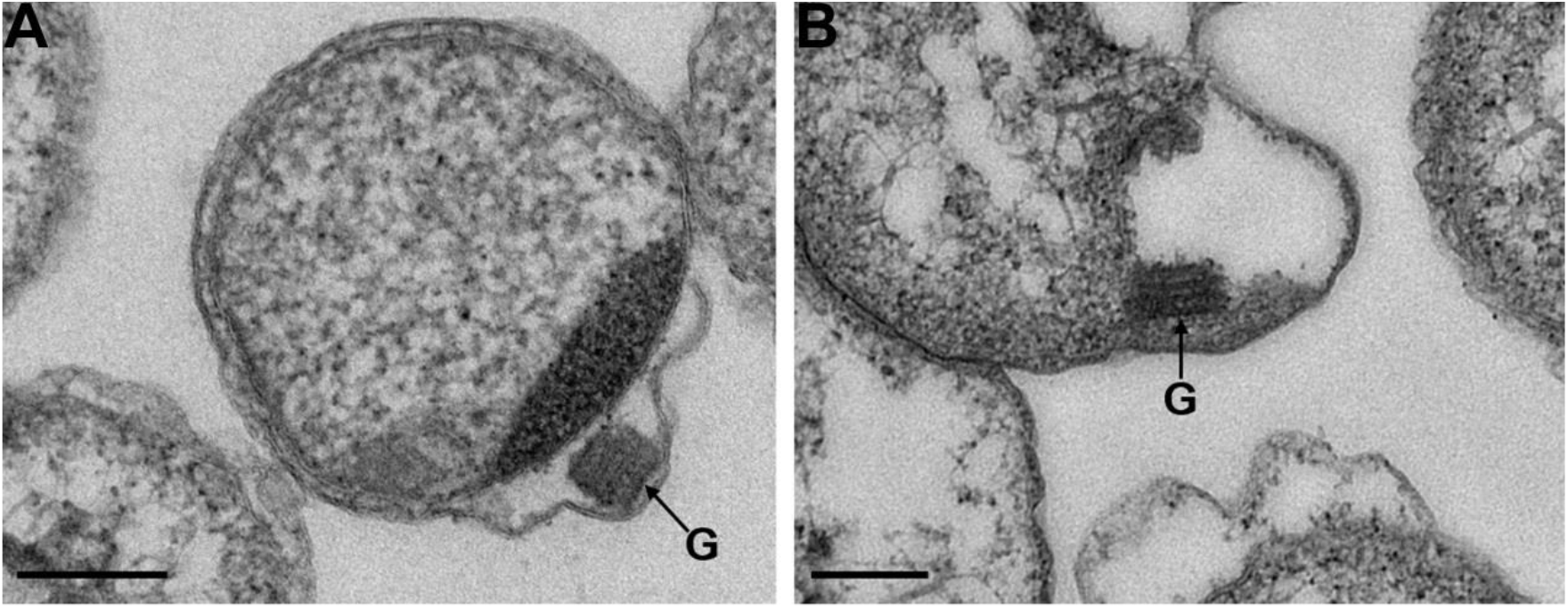
Ultrathin TEM sections showing Golgi apparatus-like organelle (indicated with G) observed in the cells of *Planctomycetes* strain ZRK32 (A-B). Bars: 200 nm.

**Expanded Data Fig. S3.**
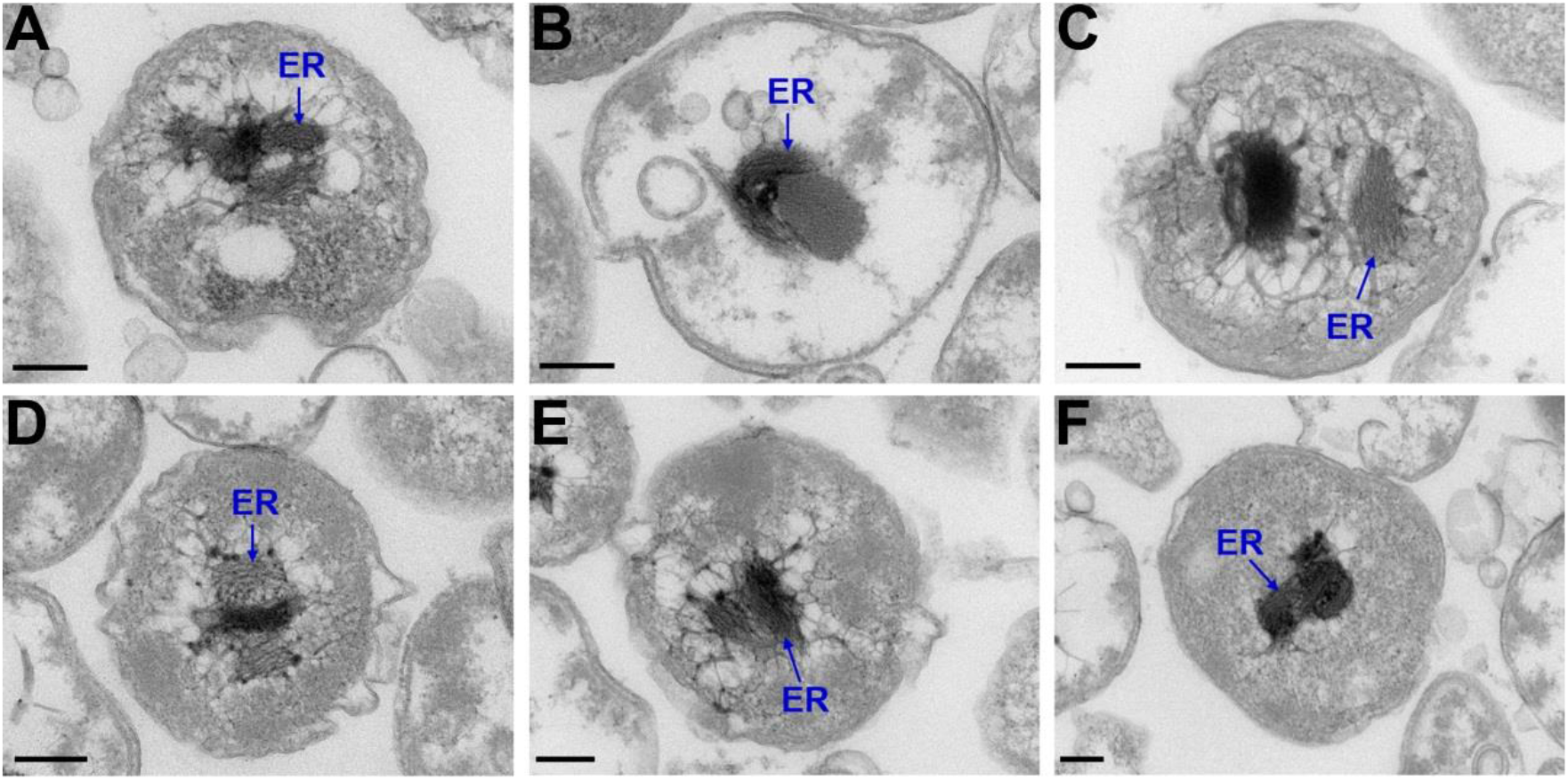
Ultrathin TEM sections showing endoplasmic reticula-like organelle (indicated with ER) observed in the cells of *Planctomycetes* strain ZRK32 (A-F). Bars: 200 nm.

**Expanded Data Fig. S4.**
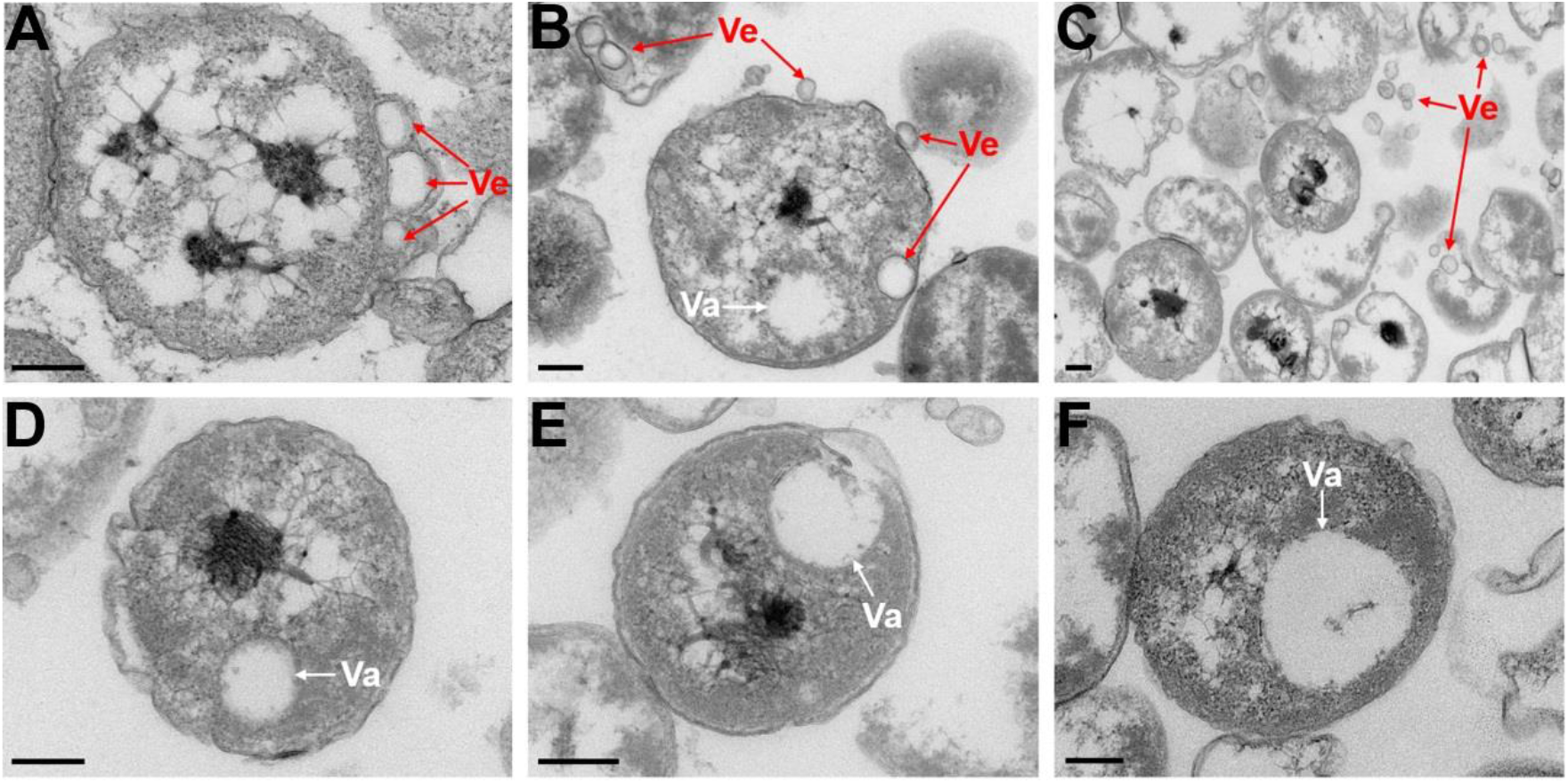
Ultrathin TEM sections showing vesicle like- (indicated with Ve in panels A-C), vacuole like- (indicated with Va in panels D-F) organelles observed in the cells of *Planctomycetes* strain ZRK32 (A-F). Bars: 200 nm.

**Expanded Data Fig. S5.**
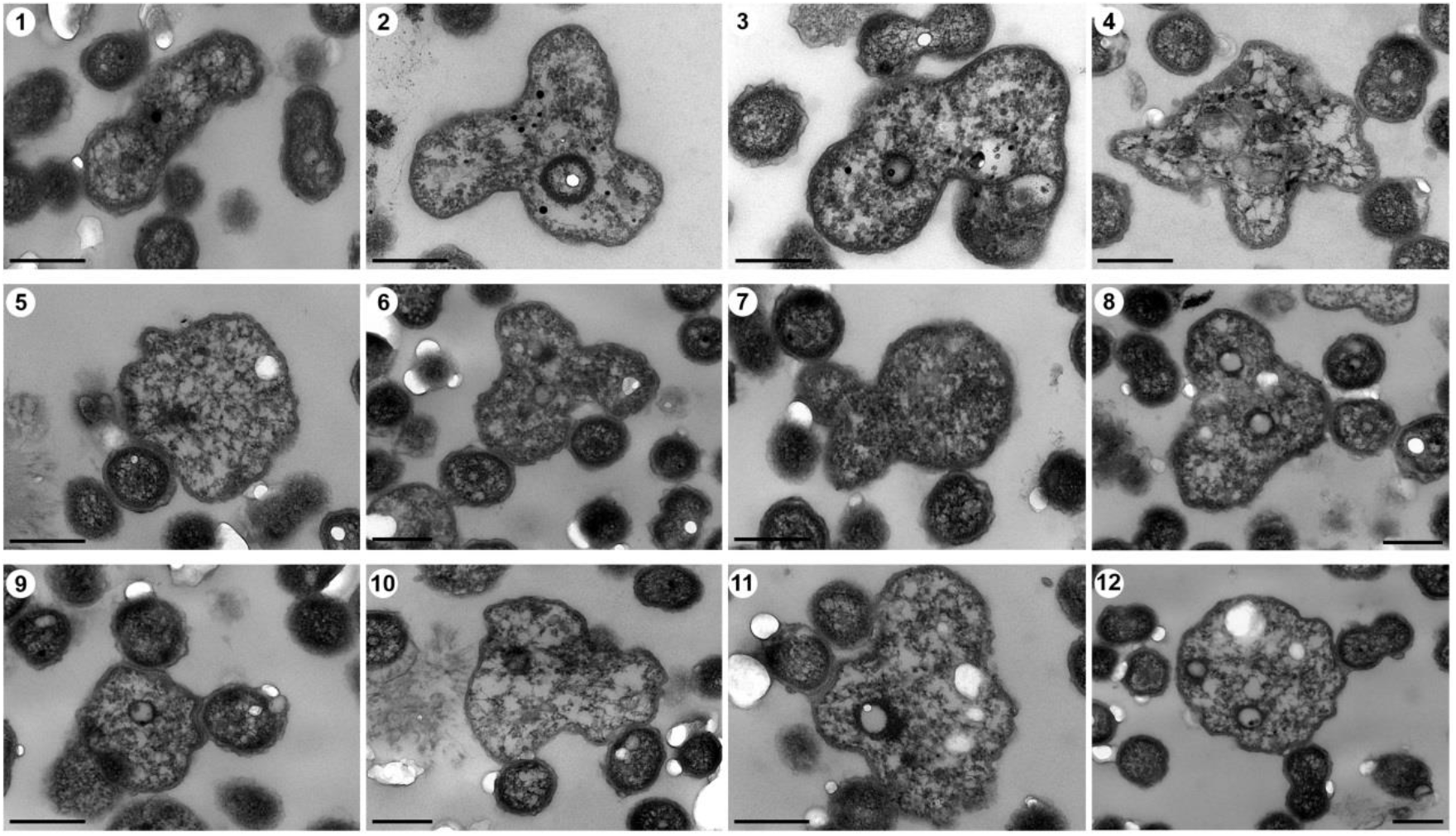
Ultrathin TEM sections showing the phagocytosis process conducted by *Verrucomicrobia* strain ZRK36.

